# Comparison of long-term outcomes between enteral nutrition via gastrostomy and total parenteral nutrition in the elderly with dysphagia: A propensity-matched cohort study

**DOI:** 10.1101/630566

**Authors:** Shigenori Masaki, Takashi Kawamoto

## Abstract

**Background:** The long-term outcomes of artificial nutrition and hydration (ANH) in the elderly with dysphagia remain uncertain. Enteral nutrition via percutaneous endoscopic gastrostomy (PEG) and total parenteral nutrition (TPN) are major methods of ANH. Although both can be a life-prolonging treatments, Japan has recently come to view PEG as representative of unnecessary life-prolonging treatment. Consequently, TPN is often chosen for ANH instead. This study aimed to compare the long-term outcomes between PEG and TPN in the elderly.

**Methods:** This single-center retrospective cohort study identified 253 elderly patients with dysphagia who received enteral nutrition via PEG (*n*=180) or TPN (*n*=73) between January 2014 and January 2017. The primary outcome was survival time. Secondary outcomes were oral intake recovery, discharge to home, and the incidence of severe pneumonia and sepsis. We performed one-to-one propensity score matching using a 0.05 caliper. The Kaplan–Meier method, log-rank test, and Cox proportional hazards model were used to analyze the survival time between groups.

**Results:** Older patients with lower nutritional states, and severe dementia were more likely to receive TPN. Propensity score matching created 55 pairs. Survival time was significantly longer in the PEG group (median, 317 vs 195 days; *P*=0.017). The hazard ratio for PEG relative to TPN was 0.60 (95% confidence interval: 0.39–0.92; *P*=0.019). There were no significant differences between the groups in oral intake recovery and discharge to home. The incidence of severe pneumonia was significantly higher in the PEG group (50.9% vs 25.5%, *P*=0.010), whereas sepsis was significantly higher in the TPN group (10.9% vs 30.9%, *P*=0.018).

**Conclusions:** PEG was associated with a significantly longer survival time, a higher incidence of severe pneumonia, and a lower incidence of sepsis compared with TPN. These results can be used in the decision-making process before initiating ANH.

## Introduction

Artificial nutrition and hydration (ANH) is a medical intervention for patients suffering from dysphagia due to various clinical conditions. ANH is administered via the enteral or intravenous route, and there are 2 representative types of ANH: Percutaneous endoscopic gastrostomy (PEG) feeding and total parenteral nutrition (TPN). PEG was initially developed as an enteral feeding technique for pediatric patients with dysphagia [1,2]. Compared to feeding via a nasogastric tube, enteral feeding via PEG can relieve laryngopharyngeal discomfort and prevent intervention failure; therefore, its use has become widespread for long-term enteral feeding in multiple patient groups including pediatric and geriatric populations [3]. However, studies have reported worse outcomes following PEG feeding in patients with dementia [4,5]; therefore, the use of PEG in elderly populations is controversial [6,7].

TPN is another common method of nutritional management [8,9]. Similar to tube feeding, TPN is also occasionally used for ANH in elderly patients with dysphagia [10]. Comparing the outcomes of enteral nutrition and parenteral nutrition are major concerns among clinicians. Previous studies have demonstrated conflicting results among patients who received enteral nutrition versus those who received parenteral nutrition [11–13].

Recently, the general population in Japan has come to view only PEG as representative of unnecessary life-prolonging treatment although both PEG and TPN can be a life-prolonging treatment. PEG is generally avoided in elderly patients; hence, a greater number of elderly patients with dysphagia choose TPN instead of PEG feeding for long-term ANH [14]. The long-term outcomes of PEG feeding versus TPN in elderly patients with dysphagia have previously been poorly documented. Therefore, we aimed to compare the long-term outcomes of PEG feeding and TPN in the elderly using propensity score-matched analysis [15–17].

## Methods

### Study design

This study was a single-center, retrospective cohort study using propensity score-matched analysis. A total of 315 consecutive elderly patients with dysphagia who underwent PEG (*n*=186) or TPN (*n*=129) for long-term ANH between January 2014 and January 2017 were considered for inclusion in the study. All PEGs were performed using the modified introducer method [18]. Central venous lines for TPN included implantable central venous ports (PORT), non-tunneled central venous catheters (NT-CVC) and peripherally inserted central catheters (PICC). We excluded patients who had advanced cancer, and those who required a PEG for gastric decompression. We also excluded TPN patients who had a PEG inserted before January 2014. Patients who received both PEG feeding and TPN between January 2014 and January 2017 were assigned to the PEG group. Finally, a total of 253 patients (180 with PEG and 73 with TPN) were included in this study.

The decision for PEG feeding or TPN was made after sufficient discussion between patients or their family and clinicians. In the TPN cases, the choices of PORT, NT-CVC and PICC were decided based on the patient’s or their family’s request and the feasibility and acceptability of each catheter in the discharge destination. Appropriate nutrition was administered based on clinical evaluation by clinicians. Clinical details were obtained from patients’ medical charts including age, gender, height, weight, underlying diseases, and blood test results. We used blood test results performed within 7 days before the start of PEG feeding or TPN. Body mass index (BMI) was calculated using the data of height and weight measured on admission. We investigated daily calorie intake on the seventh day after the procedure in both groups. We calculated the median (interquartile range; IQR) values for BMI and daily calorie intake.

Because of the anonymous nature of the data, the requirement for informed consent was waived. Study approval was obtained from the Ethical Review Board of Miyanomori Memorial Hospital.

### Outcomes

The primary outcome was defined as survival time after the start of the procedure. The secondary outcomes included oral intake recovery, discharge to home, and the incidence of severe pneumonia and sepsis. Oral intake recovery was defined as withdrawal from PEG feeding or TPN over 1 month during the observational period. Discharge to home included discharge to private residential home and housing with health and welfare services for the elderly. Definitions of oral intake recovery and discharge to home were based on that of the Ministry of Health, Labour and Welfare of Japan [19]. The diagnosis of severe pneumonia and sepsis was based on general diagnostic criteria in Japan.

### Statistical analysis

We used propensity score matching to adjust baseline differences between the groups [15–17]. The propensity score was calculated by logistic regression for estimating the probability that a patient would receive PEG feeding or TPN. We defined the following variables as potential confounders: Age, gender, underlying diseases (cerebrovascular diseases, severe dementia, neuromuscular diseases, previous history of aspiration pneumonia, ischemic heart diseases, chronic heart failure, chronic lung diseases, chronic liver diseases, chronic kidney diseases), and laboratory values (serum albumin, total lymphocyte count [TLC], total cholesterol [TC], hemoglobin and C-reactive protein) [20–26]. We performed multiple imputation to handle missing data. We created and analyzed 20 multiply imputed data sets [27,28]. The area under the receiver operating characteristic (ROC) curve was created to evaluate the performance of the logistic regression model for estimating propensity score [29]. One-to-one propensity score matching was performed to compare the primary and secondary outcomes between the groups using a 0.05 caliper, equal to 0.2 of the standard deviation of the logit of the propensity score [30,31].

We examined patient characteristics before and after propensity score matching between the groups. Continuous variables were compared with the use of the t-test or the Mann–Whitney U test, as appropriate, and categorical variables were compared with the use of Fisher’s exact test between the groups.

Survival was estimated with the Kaplan–Meier method, and the survival rate was compared using the log-rank test. We performed subgroup analysis for survival to investigate the effect of age, gender, cerebrovascular disease, severe dementia, and serum albumin. Data were censored on 28th February 2018. Cox proportional hazards models were used to estimate the hazard ratio (HR) of death for PEG feeding compared to TPN. Logistic regression analyses were used to estimate the odds ratio (OR) of outcomes. The threshold for significance was *P*<0.05. All statistical analyses were conducted using EZR version 1.37, a graphical user interface for R (The R Foundation for Statistical Computing, version 3.4.1) [32]. The Packages ‘rms version 5.1–2’ and ‘Matching version 4.9–3’ of the R software were used for multiple imputation and propensity score matching.

## Results

A total of 253 patients met the criteria for study inclusion, 180 of whom underwent PEG and 73 of whom underwent TPN. The TPN group included 28 cases of PORT, 26 cases of NT-CVC, and 19 cases of PICC. The median length of follow-up for censored cases was 601 (range, 404–823) days.

In the PEG group, missing values for TC were observed in 11 cases (6.1%). In the TPN group, missing TC and TLC values were observed in 1 case (1.4%) and 5 cases (6.8%), respectively. Missing data occurred at random because TC and TLC are not included in routine blood tests in our hospital.

Propensity score matching created 55 pairs in the PEG and TPN groups. The good fit is confirmed by the ROC curve with an area under the curve value of 0.82 (95% confidence interval [CI]: 0.76–0.87). The baseline characteristics before propensity score matching between the groups are shown in **Table 1**.

**Table 1.**
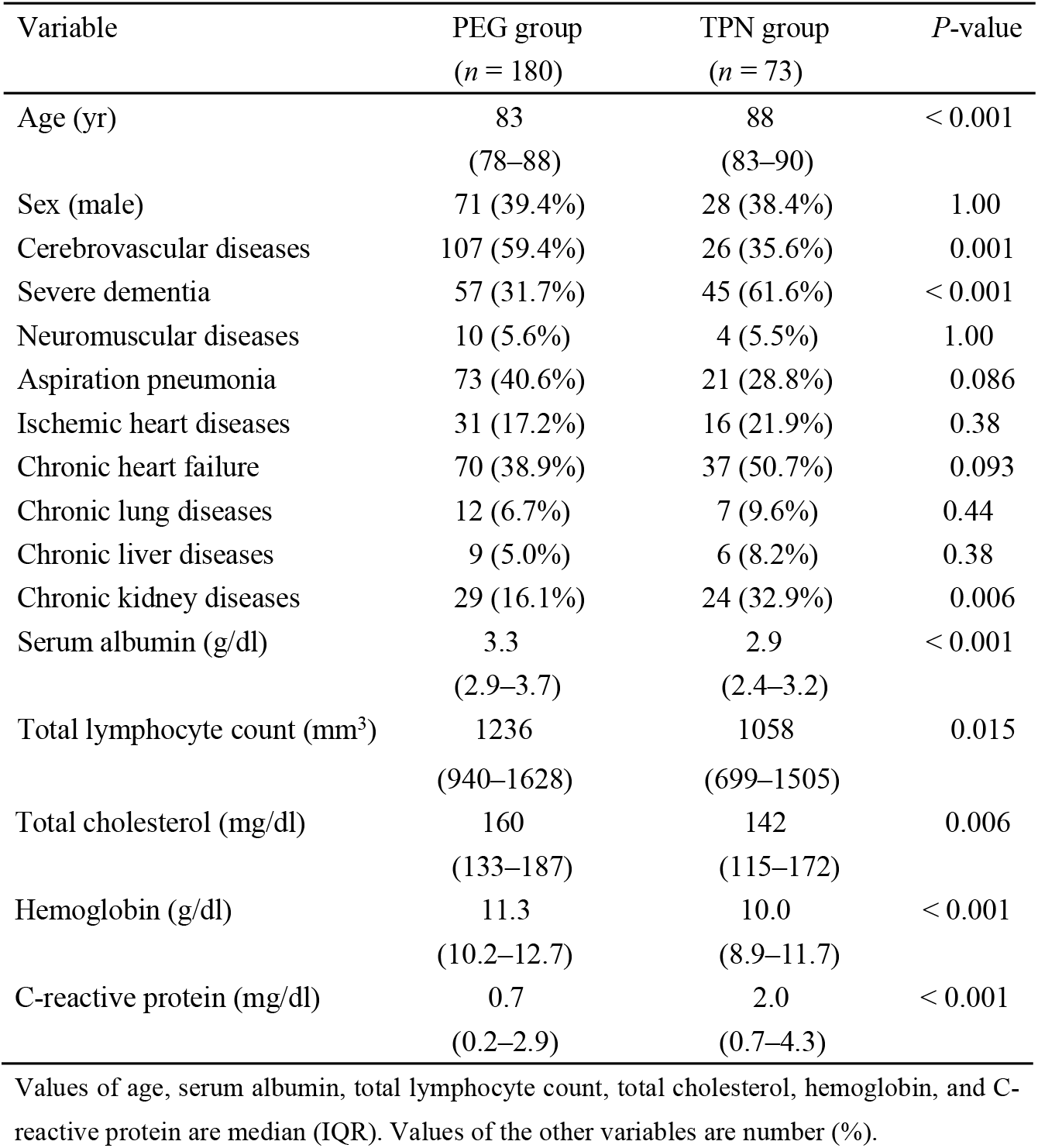
Baseline characteristics of patients before propensity score matching.

Patients with older age; severe dementia; chronic kidney disease; lower serum albumin, TLC, TC, and hemoglobin levels, as well as higher C-reactive protein levels were more likely to receive TPN. Patients with cerebrovascular disease were more likely to receive PEG feeding. The baseline characteristics after propensity-score matching between the groups are shown in **Table 2**.

**Table 2.**
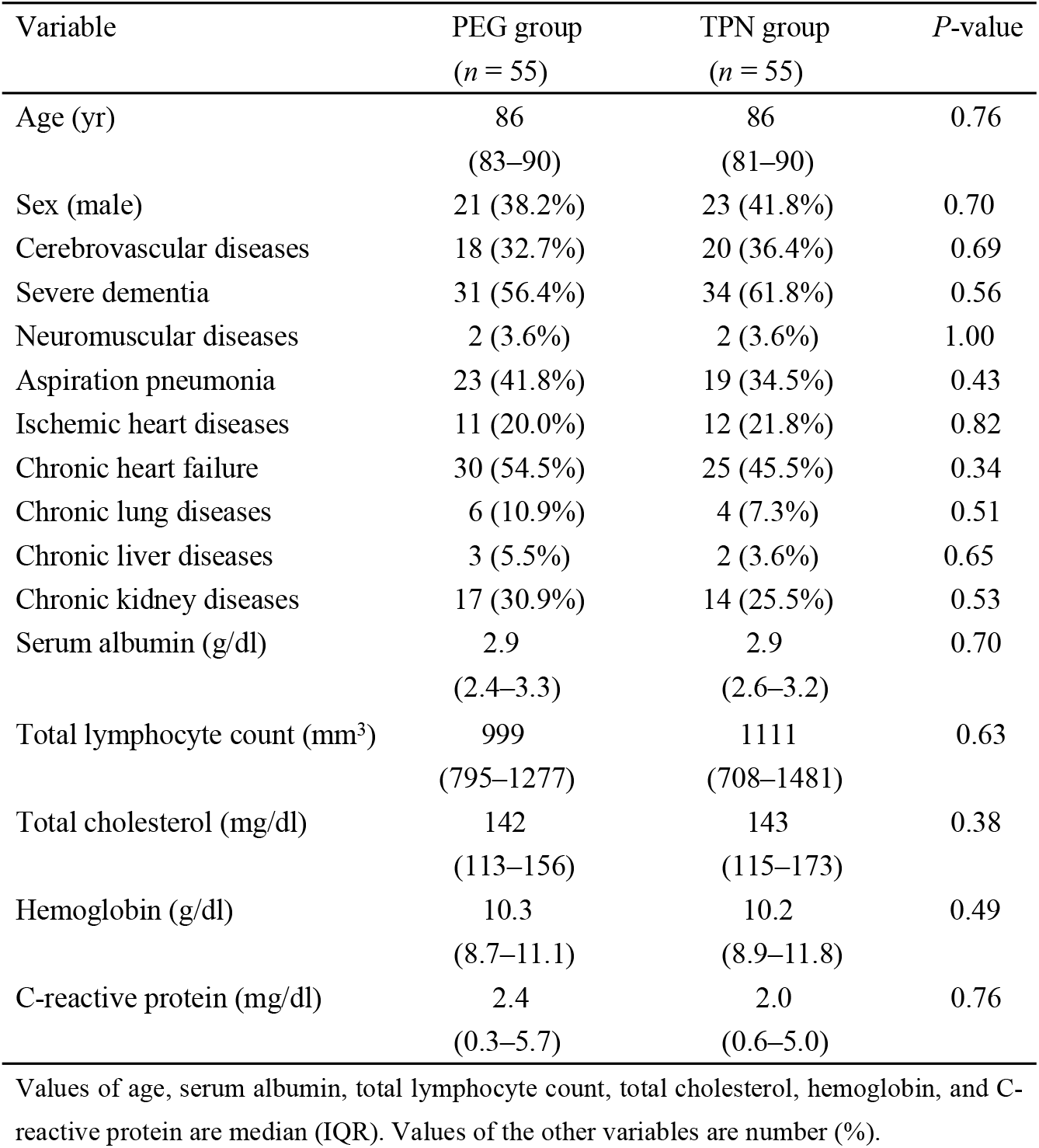
Baseline characteristics of patients after propensity score matching.

| Variable | PEG group<br>(*n* = 55) | TPN group<br>(*n* = 55) | *P*-value |
| --- | --- | --- | --- |
| Age (yr) | 86<br>(83–90) | 86<br>(81–90) | 0.76 |
| Sex (male) | 21 (38.2%) | 23 (41.8%) | 0.70 |
| Cerebrovascular diseases | 18 (32.7%) | 20 (36.4%) | 0.69 |
| Severe dementia | 31 (56.4%) | 34 (61.8%) | 0.56 |
| Neuromuscular diseases | 2 (3.6%) | 2 (3.6%) | 1.00 |
| Aspiration pneumonia | 23 (41.8%) | 19 (34.5%) | 0.43 |
| Ischemic heart diseases | 11 (20.0%) | 12 (21.8%) | 0.82 |
| Chronic heart failure | 30 (54.5%) | 25 (45.5%) | 0.34 |
| Chronic lung diseases | 6 (10.9%) | 4 (7.3%) | 0.51 |
| Chronic liver diseases | 3 (5.5%) | 2 (3.6%) | 0.65 |
| Chronic kidney diseases | 17 (30.9%) | 14 (25.5%) | 0.53 |
| Serum albumin (g/dl) | 2.9<br>(2.4–3.3) | 2.9<br>(2.6–3.2) | 0.70 |
| Total lymphocyte count (mm<sup>3</sup>) | 999<br>(795–1277) | 1111<br>(708–1481) | 0.63 |
| Total cholesterol (mg/dl) | 142<br>(113–156) | 143<br>(115–173) | 0.38 |
| Hemoglobin (g/dl) | 10.3<br>(8.7–11.1) | 10.2<br>(8.9–11.8) | 0.49 |
| C-reactive protein (mg/dl) | 2.4<br>(0.3–5.7) | 2.0<br>(0.6–5.0) | 0.76 |

Values of age, serum albumin, total lymphocyte count, total cholesterol, hemoglobin, and C-reactive protein are median (IQR). Values of the other variables are number (%).

After propensity score matching, the baseline characteristics were well balanced between the groups.

In the PEG and TPN groups, the median BMI values (IQR) were 19.0 (3.3) and 18.8 (4.8), respectively. The median daily calorie intake (IQR) was 900 (0) and 770 (250) kcal/d, respectively.

The Kaplan–Meier curve is illustrated in **Fig 1**. The log-rank test showed a significantly longer survival time in the PEG group compared with the TPN group (median, 317 vs 195 days, *P*=0.017). Cox regression analysis showed that HR for the PEG group relative to the TPN group was 0.60 (95% CI: 0.39–0.92; *P*=0.019).

**Fig 1.**
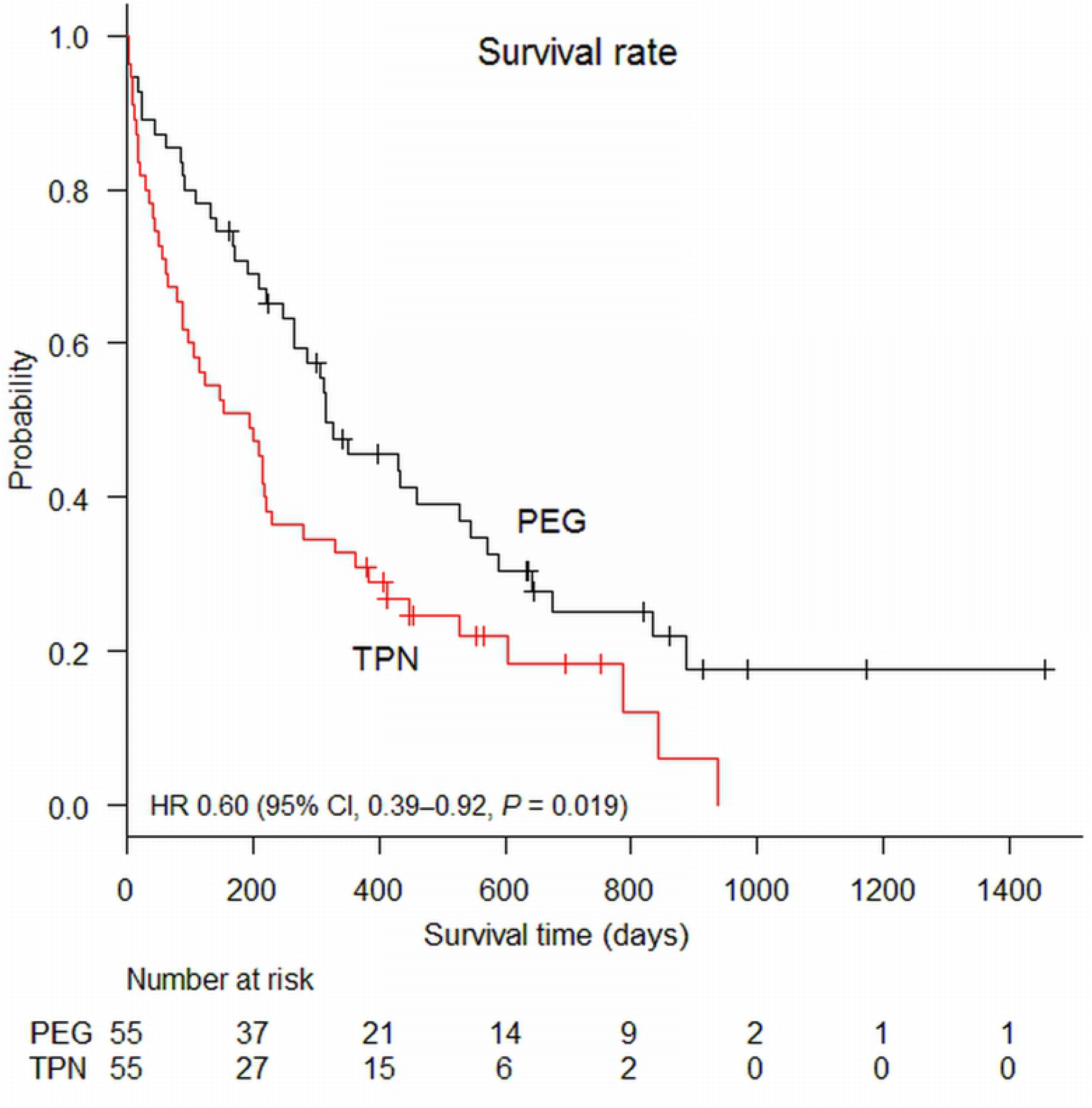
Kaplan–Meier curves of the propensity-matched groups for PEG and TPN. Propensity score matching created 55 pairs of patients. In the Cox regression analysis, HR for PEG relative to TPN was 0.60 (95% CI: 0.39-0.92; *P*=0.019).

The secondary outcomes of propensity-matched patients in the PEG and TPN groups are shown in **Table 3**.

**Table 3.** Secondary outcomes of propensity-matched patients (55 pairs) in the PEG and TPN groups.

| Outcome | PEG<br>*n* (%) | TPN<br>*n* (%) | *P*-value | <sup>a</sup> Risk difference<br>% (95% CI) |
| --- | --- | --- | --- | --- |
| Oral intake recovery | 4 (7.3) | 3 (5.5) | 1.00 | 1.8 (−7.3, +10.9) |
| Discharge to home | 7 (12.7) | 4 (7.3) | 0.53 | 5.5 (−5.7, +16.6) |
| Severe pneumonia | 28 (50.9) | 14 (25.5) | 0.010 | 25.5 (+7.9, +43.0) |
| Sepsis | 6 (10.9) | 17 (30.9) | 0.018 | −20.0 (−34.7, −5.3) |

<sup>a</sup> The risk difference for the PEG group with reference to the TPN group is shown.

There were no significant differences in the rates of oral intake recovery and discharge to home between groups. The incidence of severe pneumonia was significantly higher in the PEG group (50.9% vs 25.5%, *P*=0.010), whereas the incidence of sepsis was significantly higher in the TPN group (10.9% vs 30.9%, *P*=0.018). Logistic regression analyses of the secondary outcomes in the PEG and TPN groups are shown in **Table 4**.

**Table 4.** Logistic regression analyses of the secondary outcomes in the PEG and TPN groups.

| Outcome | <sup>a</sup> Odds Ratio (95% CI) | *P*-value |
| --- | --- | --- |
| Oral intake recovery | 1.36 (0.29–6.38) | 0.70 |
| Discharge to home | 1.86 (0.51–6.76) | 0.35 |
| Severe pneumonia | 3.04 (1.36–6.79) | 0.007 |
| Sepsis | 0.27 (0.098–0.76) | 0.013 |

<sup>a</sup> ORs for the PEG group with reference to the TPN group are shown.

ORs for the PEG group with reference to the TPN group for severe pneumonia and sepsis were 3.04 (95% CI: 1.36–6.79) and 0.27 (95% CI: 0.098–0.76), respectively. Subgroup analysis for survival is shown using a forest plot in **Fig 2**. In all subgroups, PEG consistently had a better survival compared with TPN.

**Fig 2.**
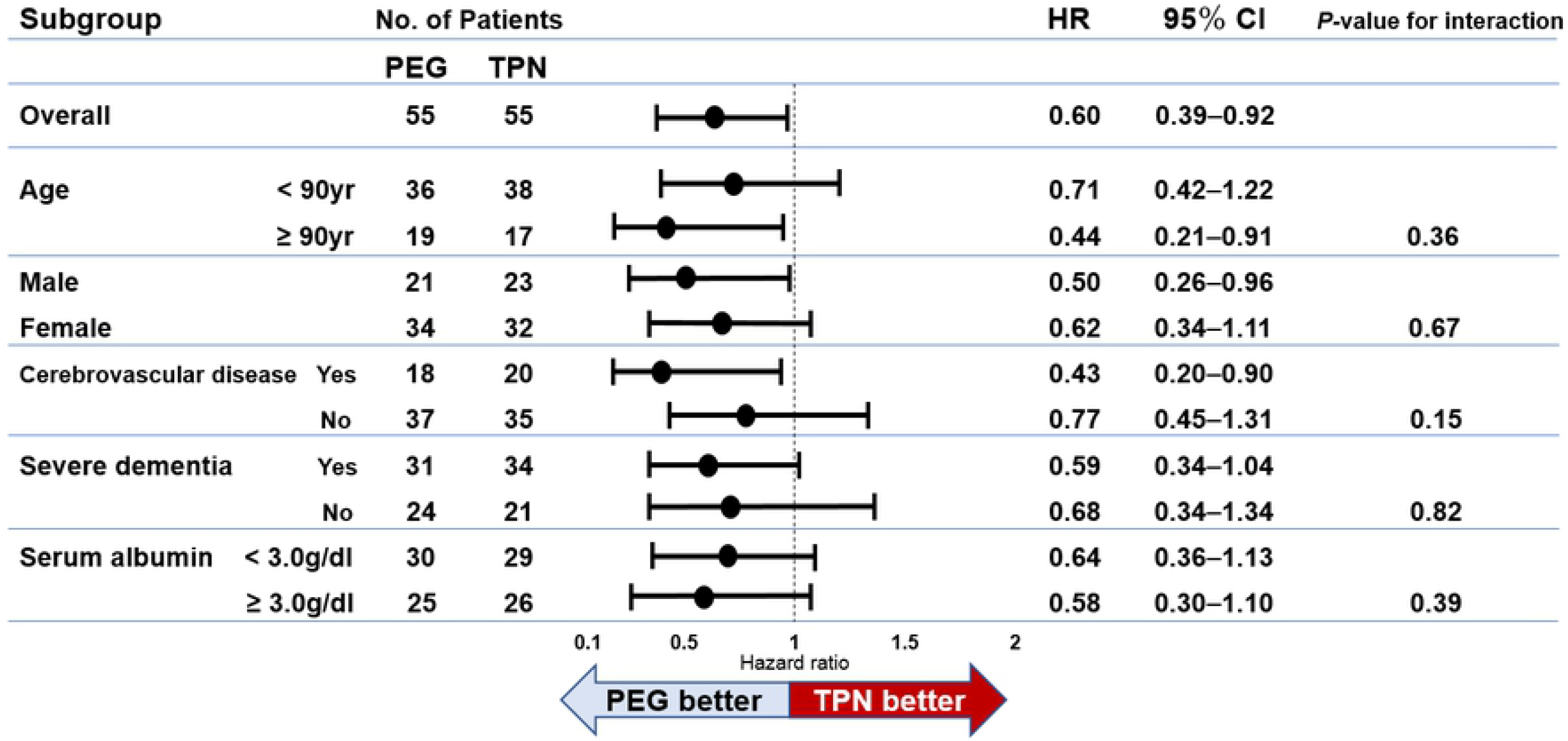
A forest plot of hazard ratios (HRs) for survival in the different subgroups. HRs from the subgroup analysis for survival between PEG and TPN are shown. HRs of < 1.00 indicate better survival in PEG compared with TPN.

## Discussion

This study investigated the long-term outcomes after PEG feeding and TPN in elderly patients using propensity score-matched analysis. We found that older patients with lower nutritional state, and severe dementia were more likely to receive TPN, whereas patients with cerebrovascular disease were more likely to receive PEG. Survival time was significantly longer in the PEG group. The incidence of severe pneumonia was significantly higher in the PEG group whereas that of sepsis was significantly higher in the TPN group.

Previous studies that compared the outcomes of patients managed with enteral nutrition and parenteral nutrition demonstrated conflicting results. For example, with respect to mortality, studies found that enteral nutrition was associated with lower mortality rates [11] or no effect on overall mortality [33]. It has also been demonstrated that enteral nutrition is associated with a lower risk of infection [33,34], a higher rate of postoperative complications rate, and a lower rate of early recovery of oral feeding after operation [13] compared to parenteral nutrition. The general rule is that enteral feeding should be considered in patients with normal digestive function whereas TPN should be used if enteral nutrition is not feasible [35]. Contrastingly, ANH for elderly patients with dysphagia can be a life-prolonging treatment [36]; therefore, the choice of enteral versus parenteral nutrition is not only based on the digestive function of the patients but also on their clinical condition and the preferences of the patients and their family members [14,36,37]. This may result in selection bias and differences in the baseline characteristics of the PEG feeding and TPN study groups; therefore, we performed propensity score matching to adjust baseline characteristics to compare the effect of PEG feeding and TPN more accurately [15–17,23].

In this study, a comparison of baseline characteristics between the groups before propensity score matching revealed that patients with older age, lower serum albumin levels, higher C-reactive protein levels, and severe dementia were more likely to receive TPN. Older age, lower serum albumin levels, higher C-reactive protein levels, and severe dementia were reported as poor prognostic factors after PEG [4,5,20-22,38]. Our results indicated that PEG tended to be avoided in patients with such poor prognostic factors, and as a result, TPN was chosen as the alternative modality for ANH. Furthermore, TLC, TC, and hemoglobin were significantly lower in the TPN group than in the PEG group before propensity score matching, suggesting that TPN tended to be chosen for patients with a poorer general condition.

Survival analysis showed better results in the PEG group than in the TPN group. This may be explained by the fact that enteral nutrition has gastrointestinal, immune, and metabolic benefits compared with parenteral nutrition [35,39,40]. Additionally, in this study, the daily calorie intake was higher in the PEG group than in the TPN group. This difference between groups may have affected the results of the survival analysis. Previous studies showed that PEG did not improve survival in patients with dementia [4,5,38]. On the contrary, it has been reported that dementia was not a significant prognostic factor after PEG [41]. In our subgroup analysis, PEG was associated with better survival than TPN even in patients with severe dementia. Furthermore, compared to TPN, PEG showed a survival benefit regardless of age, sex, cerebrovascular disease, and serum albumin level. These results suggested that enteral nutrition still had a better impact on survival even in elderly individuals with a poorer general condition.

Most of the previous studies that compared enteral and parenteral nutrition defined survival and infection rates as the primary and secondary outcomes, respectively [11,13,23,25,33,34]. Here, we placed importance on quality of life after the start of ANH, and thus we chose oral intake recovery and discharge to home as the secondary outcomes. Previous studies showed that age and BMI were predictive factors of oral intake recovery in stroke patients with tube-feeding [42,43]. In this study, age and BMI were similar between groups, and there were no significant differences in oral intake recovery between groups. Oral intake recovery rates were low in both groups, with most patients requiring continuous ANH. Moreover, there were no significant differences in discharge to home between groups, indicating that both PEG feeding and TPN were feasible in a home environment [9,35,44,45]. However, the proportion of patients being discharged to their homes was also not high in either group, suggesting that most of the elderly patients with dysphagia requiring ANH were bound to stay in long-term care facilities rather than their own homes regardless of receiving PEG feeding or TPN. It is necessary to provide patients and their family members with information regarding the general clinical course to aid their decision-making process before initiating ANH [46]; our results add to such clinical information for supporting the decision-making process.

The incidence of severe pneumonia was significantly higher in the PEG group. This result was expected and clinically plausible because enteral nutrition administered via PEG poses a risk of gastroesophageal reflux and aspiration pneumonia owing to the underlying pharyngeal and laryngeal dysfunction of patients who require feeding through this modality [23,35,47]. Switching from PEG feeding to TPN may be an option for patients who underwent PEG feeding and repeatedly suffered from aspiration pneumonia because TPN is more effective in reducing the risk of severe pneumonia than PEG feeding. In contrast, as expected, the incidence of sepsis in the TPN group was significantly higher than that in the PEG group. This may be due to the fact that TPN has been associated with catheter-related bloodstream infections and bacterial translocation [34,48–50]. Furthermore, the use of NT-CVC for long-term TPN may affect the rate of catheter-related bloodstream infections and the incidence of sepsis in the TPN group [51].

Several limitations of this study should be acknowledged. First, this was a retrospective observational study without randomization; therefore, assignment to each group may have been biased. Although propensity score matching was used to adjust the differences in baseline characteristics, the results may still have been biased because of unmeasured confounders. Second, the results of this study are applicable only to these patients who were included in the paired analysis, and therefore the results may not be generalizable to a broader population. Third, certain patients in the PEG group received not only PEG feeding but also TPN depending on their clinical condition, and furthermore, the daily calorie intake was not equal between the groups. Fourth, this was a single-center study with a small sample size.

## Conclusions

In summary, we performed a propensity-matched analysis to compare the outcomes of PEG and TPN in the elderly. We found that compared to TPN, PEG was associated with better survival and a higher incidence of severe pneumonia as well as a lower incidence of sepsis, with no significant inter-group differences noted in oral intake recovery and discharge to home. Further studies with a larger sample size and randomized controlled design are required.

## Acknowledgments

We would like to thank Editage (www.editage.jp) for English language editing.

